# Bacmeta: simulation for genomic evolution in bacterial metapopulations

**DOI:** 10.1101/175257

**Authors:** Aleksi Sipola, Pekka Marttinen, Jukka Corander

## Abstract

The advent of genomic data from densely sampled bacterial populations has created a need for flexible simulators by which models and hypotheses can be efficiently investigated in the light of empirical observations. Bacmeta provides fast stochastic simulation of neutral evolution within a large collection of interconnected bacterial populations with completely adjustable connectivity network. Stochastic events of mutations, recombinations, insertions/deletions, migrations and microepidemics can be simulated in discrete non-overlapping generations with a Wright-Fisher model that operates on explicit sequence data of any desired genome length. Each model component, including locus, bacterial strain, population, and ultimately the whole metapopulation, is efficiently simulated using C++ objects, and detailed metadata from each level of the simulation can be acquired. The software can be executed in a cluster environment using simple textual input files, enabling, e.g., large-scale simulations and likelihood-free inference. Bacmeta is implemented with C++ for Linux, Mac and Windows. It is available at https://bitbucket.org/aleksisipola/bacmeta under the BSD 3-clause license.

**Supplementary information:** Supplementary data are available online at bioRxiv.

## 1 Introduction

Simulation models can be used for prediction, parameter estimation, and for validating methods used in population genomics (Hoban *et al.*, 2012). Most general-purpose simulators are tailored mainly for eukaryotes (e.g., Arenas and Posada, 2014). However, many studies on evolutionary processes in bacteria have emerged recently, using simulation software tailored for their specific purposes (Fraser *et al.*, 2007; Friedman *et al.*, 2013; Marttinen *et al.*, 2015; Niehus *et al.*, 2015; Numminen *et al.*, 2016). Simulators can be divided into two categories (Hoban *et al.*, 2012): coalescent simulation starts with the present-day population and simulates backwards in time, coalescing individuals until the most recent common ancestor is found, while forward simulators maintain a population of individuals and simulate forward in time by sampling the next generation from the current one. In general, coalescent simulators are faster, by only considering the ancestors of the current individuals, but forward simulation allows greater flexibility to define the model. This makes the latter particularly attractive for bacteria, where recombination shuffles genetic material between genomes in a complex manner that depends, for example, on the genetic and physical distance between the donor and recipient strains. Furthermore, recombination may cause different parts of the genome to have completely distinct population histories (Feil *et al.*, 2001; Mostowy *et al.*, 2017), undermining the assumption of a single coalescent. The recently published general-purpose simulators tailored for bacteria have all been based on the coalescent approach (Brown *et al.*, 2016; De Maio and Wilson, 2017). Hence, there is a need for an efficient general-purpose forward simulator for bacterial population genomics.

Bacmeta provides an efficient C++ implementation of a finite metapopulation Wright-Fisher model with explicit genome sequences evolving for each strain present in the metapopulation. Use of shared pointers of C++ and compact object representations result in low memory and runtime requirements. The model allows multiple arbitrarily connected populations, each with thousands of bacteria, for which the genome sequences are subjected to evolutionary events over discrete non-overlapping generations. Bacmeta implements a large variety of different event types governed by user-defined parameters using simple textual input files, which provides a convenient framework for large-scale simulations, integration with other software and likelihood-free inference. For example, Bacmeta could be used for testing methods for inferring recombination (Croucher *et al.*, 2014; Didelot and Wilson, 2015; Mostowy *et al.*, 2017), since every past evolutionary event can be stored to provide the ground-truth. Another potential application is likelihood-free inference for model parameters, as in Marttinen *et al.* (2015); Numminen *et al.* (2016); De Maio and Wilson (2017), based on the Approximate Bayesian Computation inference framework (Beaumont *et al.*, 2002; Lintusaari *et al.*, 2017).

## 2 Features

The main input parameters for Bacmeta are defined in a plain ascii text file. Optional migration parameters can be given in a second input file. Examples of these are given in Tables S1 and S2. The simulation of each evolutionary event, including reproduction, is executed at each generation for a desired number of iterations. The events that can be simulated are displayed in Figure 1A and B.

**Figure 1:**
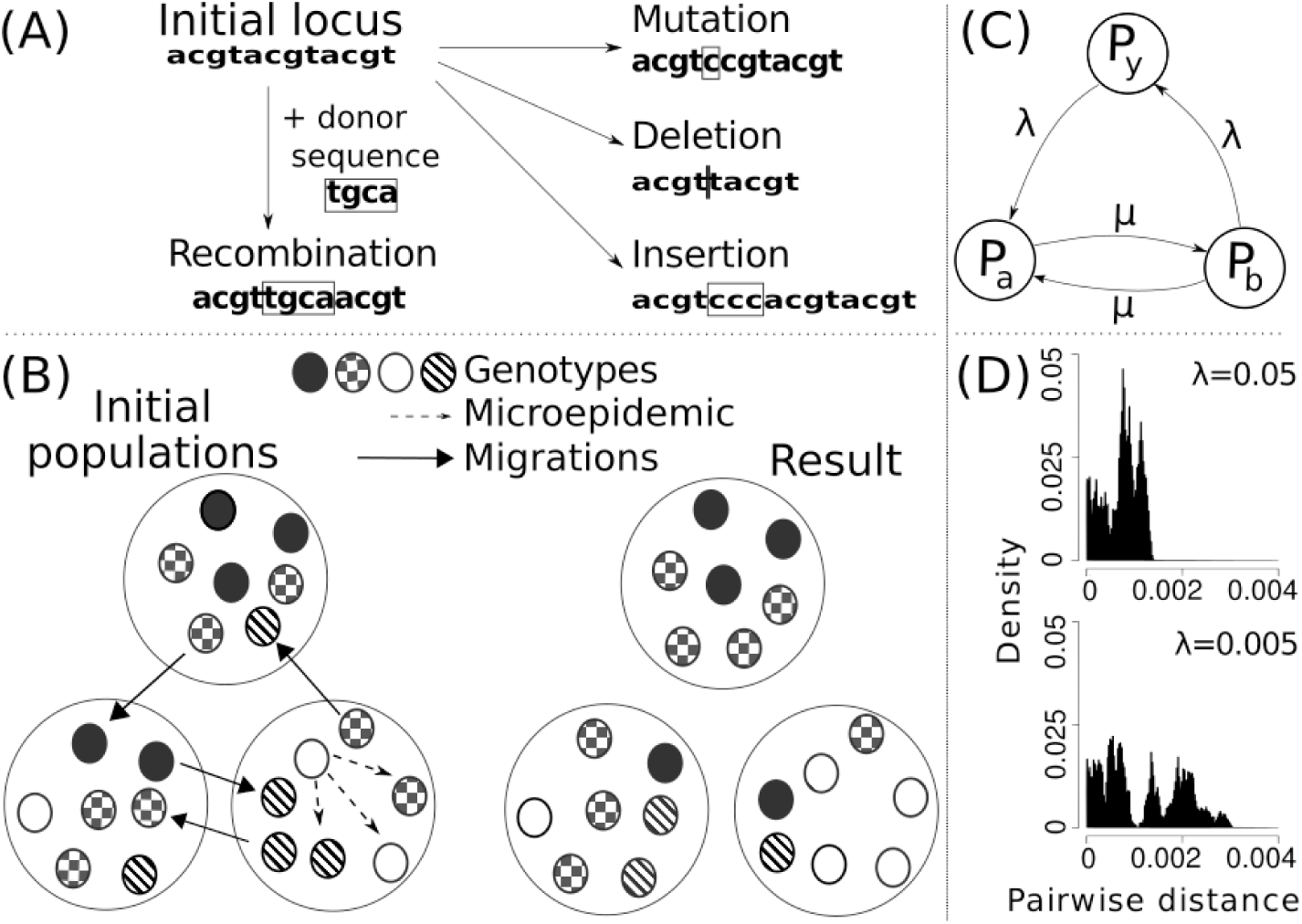
(A) Illustrations of the evolutionary events and (B) illustrations of population dynamic events, excluding random sampling between generations. (C) Example case: The migration connectedness of metapopulation as a network graph, where *Py* is the observed population and edge weight *λ* represents the inward and outward migration rates of *P*_*y*_ and *μ* = 0.01. (D) Effect of low versus high value of *λ* on pairwise distances in population *P*_*y*_. Computed from 10 observed simulations per *λ*-value.

The order of the events can be fixed or random. Each generation ends in the selection of bacteria for seeding the next generation by random sampling with replacement. The count of each event type per generation is modeled as a Poisson process with a user-defined rate parameter. For migration and microepidemic events we use the parametrization introduced by Numminen *et al.* (2016). For mutations and recombinations we use the same approach as Marttinen *et al.* (2015), except that mutations are generated under an explicit mutation model with separate user-defined weights for all nucleotide pairs in ACGT. For insertion/deletion sizes we use the model presented by Benner *et al.* (1993), with rate defined in relation to mutations. Note that this also allows for more coarse-grained summaries of the produced data, for example as an infinite alleles type integer labeling for each locus, useful for certain types of genomic analyses. Haplotypes can be flexibly represented as genomic islands of any desired length and number, such that separate genomic regions or secondary chromosomes can be imitated. Outputs from the simulator include synthetic DNA sequences, pairwise distance measures and several different summaries, e.g., counts of the different events.

## 3 Example case

For an illustrative example of the functionality and performance of Bacmeta, we considered the effect of inter-population connectedness via migration parameter setup. We simulated 10 occurrences of the cases: *i*) low connectedness of observed population *P*_*y*_, and case *ii*) high connectedness of observed population *P*_*y*_, each for 20 000 generations. We used a metapopulation consisting of three populations, with migration routes as illustrated in Figure 1C and the migration rate input file displayed in Table S2. General simulation parameters were as given in Table S1 and followed values of recombinogenic bacteria. Figure 1D shows results. As expected in this setup, the higher connectivity led to markedly lower pairwise distances, due to the reduced divergence between the populations. Runtimes for these simulations were between 35-50 seconds on a single core of Intel Core i5-7200U CPU @ 2.50GHz.

